# Evolutionary Lags in the Primate Brain Size/Body Size Relationship

**DOI:** 10.1101/2024.02.05.578865

**Authors:** Robin Dunbar

## Abstract

**INTRODUCTION:** The original brain lag hypothesis proposed that primate brain evolution depended on spare energy derivative of savings of scale enabled by increasing body size. Deaner & Nunn [1] concluded that, in fact, there was no evidence for a brain lag. However, their result may have been due to a number of possible confounds in their analysis.

**METHODS:** I revisit their analysis to test for potential confounds using updated datasets. I also ask how primates paid for the energy costs incurred by changes in brain and body mass, and whether the impetus for these changes was predation risk. Finally, I ask whether the observed patterns explain the brain/body size ratio trajectory observed in fossil hominins.

**RESULTS:** I show that using statistically more appropriate statistics and updated data yields a significant brain lag effect. However, contrary to the original brain lag hypothesis, the brain/body ratio does not converge back on the allometric regression line, but continues to evolve beyond it. Increases in brain size are correlated with exploiting large group size rather than body size as the principal defence against predation risk, with significant growth in brain size (but not body size) only being possible if species adopted a more frugivorous diet. Finally, I show that hominins followed a similar trajectory from an australopithecine baseline that fell on the relevant allometric regression.

**CONCLUSION:** The brain lag effect is much more complicated than the original hypothesis proposed, with a distinctive switch from body to brain over evolutionary time.

## Introduction

Ever since Jerison’s [2] seminal analyses, it has been known that, across mammals in general and primates in particular, brain size is correlated with body mass in an allometric power relationship, with different orders clearly forming distinct grades. An early suggestion was that much of the internal variance around the order-specific allometric regression line, especially in large-brained primates, is a consequence of a lag effect in which body size changes first, with brain mass taking time to catch up with the changes in body mass [2–8].

This assumption was based mainly on the fact that body size is relatively labile, and can vary considerably even within species as a function of local environmental conditions [9–10], whereas the complex interconnectivity of brain systems means that it is likely to take longer to bring together the necessary genetic changes without disrupting functional neural systems. The fact that changes in body size clearly outstripped changes in brain size during domestication [11–13] (notwithstanding the rather questionable claims to the contrary by [14]) lends support to this. Given this, it was assumed that, given enough time, brain and body size would converge back on the common allometric relationship as brain size expanded to absorb the additional energy made available by a larger body mass through the efficiencies of provided by Kleiber’s Law [15–17].

Deaner & Nunn [1] developed a particularly innovative method for testing the brain lag hypothesis that involved plotting the residuals of the phylogenetic contrasts in brain mass regressed on the contrasts in body mass against estimated date of divergence. Since the original hypothesis assumed that body mass will always change first, the lag would will necessarily be directional. Hence, they argued, the magnitude of the signed residual from the brain-to-body size allometric relationship should be positively correlated with divergence date: when body mass contrasts are constrained to be positive, ‘young’ nodes (i.e. recent speciation events) should consist of species pairs with strongly negative residuals (smaller brains relative to body size), whereas nodes with deeper divergence dates (that have had more time to readjust) will eventually approximate the allometric line (where residuals equal 0). They found that, for a sample of primates, there was no correlation between the two variables, neither for males (r=-0.20, p=0.38) nor for females (r=0.15, p=0.480), or when controlling for circadian rhythm. This led them to conclude that there was no evidence for the lag hypothesis. By implication, brain and body size must in fact change simultaneously under the same environmental pressure, just as is the case for brain size and group size in primates (but not other mammals) [18]. Because these results seemed conclusive, the hypothesis lost traction and was no longer discussed in the literature.

More recently, however, a number of separate developments in our understanding of primate biology and evolution have raised the possibility that the original Deaner-Nunn result by be a consequence of several potential sources of confound. I identify four such issues that that were not appreciated at the time of the original analysis.

One is the fact that the slope in the mammalian brain/body size regression differs across the different orders [19–21], and even between families within an order [22]. With the exception of Shultz & Dunbar [19], these studies have not, however, offered any explanation as to why or how these grades might have evolved. Rather, they essentially show that rate changes in the brain/body slope can be reliably identified. If the primate brain/body dataset is not an homogenous set, but consists of grades in the way the primate social brain dataset does [23–26], then the Deaner-Nunn analysis falls foul of the Yule-Simpson effect (a version of the Ecological Fallacy) which will lower the slope in OLS regressions [27,28]. This problem that was discussed at some length during the early days of comparative analyses [29–32], but has largely been forgotten about since. Partitioning the data by grades can yield very different results [28].

Second, their analysis assumed that brain size is largely a function of body size, with the two being genetically linked. But if brain size and body size are genetically uncoupled, as they appear to be in primates [3], there is no principled reason why body size should always have to change first or why brain size should subsequently have to “catch up” with body size. That brain and body mass are not yoked in close genetic linkage is confirmed by breeding experiments showing that they can undergo independent selection, at least in the short term [33]. Lande [3] examined the brain-to-body size relationship across mammals in order to evaluate the evolutionary coupling between these traits. He argued that the genetic correlation between brain and body size in primates is unusually weak compared to other mammalian orders. More importantly, he found that the variance in the primate allometric relationship increases with body size, suggesting that the two become progressively decoupled as body size increases [20,34–35].

Third, although Deaner & Nunn reported that there was no correlation between their index of residual brain size and social group size, one plausible explanation for the variability in the residuals of brain size regressed on body size is that they in fact reflect different species pursuing alternative anti-predator strategies. Large body size and large group size have long been known to be alternative strategies that primates use (sometimes even in combination) to reduce predation risk in order to occupy habitats subject to high predator densities [36–39]. In primates at least, group size has been shown to correlate strongly with brain size, both within (humans [40–47]; monkeys [48–50]) and between species [18,19,24–26, 28, 51–53], because the cognitive demands imposed by increasing social group size in the kind of bonded societies characteristic of primates are neurologically very demanding [25,54–57]. (Note: the few cases claiming that brain size and group size are uncorrelated all turn out to be statistically flawed: they inadvertently tested hypotheses about constraints, not hypotheses about selection effects. For details see [28].)

The fourth issue is simply the fact that the datings on our phylogenies have changed significantly in the last three decades as a result of improved molecular genetic data. As a result, the datings for many of the last common ancestors of allied lineages have changed dramatically. Improved datings, especially for the deeper common ancestors, may change the results completely.

The original brain lag hypothesis assumed that body size changed first (causing initial residuals of brain on body mass to be negative). Deaner & Nunn therefore tested for a positive relationship between brain/body residuals and time since divergence. Strictly speaking (though they never tested for this), if the brain lag hypothesis is true, then residuals should converge back onto the allometric relationship and remain there until further deflected. In effect, the residuals should oscillate around the allometric regression line. Hence, if the residuals systematically project beyond the allometric regression (and do so more with time since divergence), this would be evidence against the lag hypothesis as originally conceived. If this was the case, however, it would open up the rather more interesting possibility that a brain lag might allow species to catapult themselves onto a higher brain/body grade by exploiting the opportunities offered by Kleiber’s Law: larger body size creates greater savings of scale in metabolic energy demand, thereby allowing the surplus energy intake to be diverted into evolving an even larger brain.

Here, I reanalyse the Deaner-Nunn data to determine whether a brain lag is evident if (a) we use updated phylogenetic divergence times and (b) we take grades into account. If there is evidence for a brain lag, this then raises several further questions about the direction of the lag, why the lag occurs (i.e. the environmental factors that select for a particular change) and how species balance the energy demands inevitably created by any such change. I test the hypothesis that deviations from the allometric regression line are a response to high predation risk as the known principal driver of primate social evolution [25,38,40,52,59] and, then, whether changes in brain or body size necessitated dietary changes to meet the energetic costs of the additional tissue. Finally, I use the excellent fossil record for hominins to test the evolutionary trajectory suggested by these analyses.

## Methods

Deaner & Nunn’s approach to the problem of how to test the brain lag hypothesis was particularly ingenious. They suggested using standard CAIC contrasts to identify residuals from the common allometric regression and plotting these against divergence time. They predicted that when body mass is constrained to change first, a regression of residuals against time since divergence would yield a positive slope if the lag hypothesis was true. For these purposes, contrasts were measured without reference to elapsed time since the last common ancestor (the Brownian motion control) specifically in order to be able to use divergence time as the independent variable in the analysis. In addition, they explicitly used only nodes that were tip comparisons (i.e. comparisons between living species) and avoided ancestral nodes at higher levels in the phylogeny since these can never be known with any precision. Even small degrees of error variance in the measurements at deep nodes risks destabilising the statistical analysis by introducing too much unnecessary error variance (thereby lowering the slope of any OLS regression).

If we are to test the validity of the original analysis, it is essential to use exactly the same dataset and statistical methods as Deaner & Nunn [1] did in order to be sure that any differences in findings are due to the analysis and not to differences in the data or methods. Deaner & Nunn also insisted that brain and body size estimates must be from the same individual, otherwise we risk confounding phenotypic environmental effects with evolutionary effects. They therefore rejected the larger samples then available from studies that used cranial volumes (ECV) to estimate brain size because these are invariably based on museum specimens sourced from a variety of locations, with body masses derived from a mixture of field and zoo studies. For this reason, they deliberately restricted their analysis to the Stephan et al. [60] dataset where brain and body weights were determined from the same anatomical specimen. Although more comprehensive larger datasets have subsequently become available for a wider range of species [61–63], these all use endocranial volumes rather than actual brain mass or mix ECV with actual brain volume data. Rilling [64] strongly cautioned against mixing brain data based on different methods because of the different error variances involved.

In addition, Deaner & Nunn specifically cautioned against using ECVs as a proxy for brain size because ECV includes intracranial spaces not occupied by actual brain tissue (the subdural spaces and ventricles); as a result, ECV estimates are typically ∼5-10% larger than actual brain size [65] with the difference being larger in large-brained species [2]. This will clearly change the brain:body mass ratio, especially during the initial and late timescales when sister taxa are, respectively, just beginning to diverge or are converging back on the allometric line. I therefore follow Deaner & Nunn in using the Stephan dataset since only these data are based on brain and body weights taken from the same individual. Although this reduces the sample size, in fact the reduction is modest. Using the Boddy et al. [62] dataset would yield data for only two additional taxa that are not already represented in the Stephan dataset. One of these is based on ECV, and the other uses brain and body mass values from different populations. In effect, no additional contrasts would in fact be available even if we accept ECVs.

The Stephan et al. primate brain dataset yields 31 tip nodes for which contrasts in brain mass and contrasts in body mass are available for matched pairs of species. The species and contrasts as used by Deaner & Nunn are listed in Table S1, distinguishing those that are within-genus from those that are from different genera (between-genera). Since contrasts at different phylogenetic depths vary considerably in timing, this allows a wide range of divergence dates to be considered. This was important both because we do not know how long it might take for sister taxa to converge back on the common allometric regression line. Within-genus contrasts represent more recent divergences (typically with divergence dates <10 Ma), which allow us to identify which variable typically changes first.

Deaner & Nunn used the divergence dates given by Purvis [66]. While undoubtedly the best available at the time, these were based largely on anatomical traits and fossil dates. Since then, more accurate molecular genetic phylogenies have become available, and I here use the genetic divergence times provided by Perelman et al. [67]. Although the two divergence estimates correlate significantly (r=0.732, N=25, p<0.0001), the agreement on some comparisons between taxonomically very different species (e.g. lemurines versus Old World monkeys) is poor. I ran the analysis with both sets of divergence dates in order to determine whether this was a potential source of confound.

Finally, I ask whether the findings from this analysis help us to understand hominin brain evolution. Largely for taphonomic reasons, brain evolution within the hominin lineage is unusually well documented compared to other primates. However, in this case, we necessarily have to rely on ECV data as an index of brain mass. I use the ECV volumes and earliest appearance dates from [68], and species body sizes from [69] based on the same specimens. Hominin ECV values are reduced by 5.9% (following [70]) to give an improved estimate of actual brain mass. Body mass data for chimpanzees are from [71]; wild chimpanzee brain data are from [72]. In this case, we are interested in the changes that have occurred since the hominin/*Pan* split (∼7.0 Ma), and hence all contrasts are between individual hominin species and living chimpanzees (*Pan*). As the time base, I calculated the time elapsed from the divergence of the hominin and chimpanzee lineage at ∼7 Ma to the appearance of the earliest member of each subsequent species in the hominin lineage. Dates for the last common ancestor vary between 6-8 Ma, but the difference is of little importance because we use the same reference point for all contrasts. Errors in the dating will simply move the distribution of points to the left or right on the X-axis, and only by at most a matter of a million years. This will have only a limited effect on any regression parameter estimates.

Having determined whether or not there is a brain lag effect, I ask two further important questions: (1) what selection forces drove the deviation from the allometric relationship and (2) how do the species concerned deal with the additional energetic demands of larger brains and larger bodies. Deaner & Nunn tested whether the residuals in the brain/body size relationship predicted social group size as a potential selection pressure for deviating from the allometric regression line. However, primates exploit both increased body mass and increased group size as solutions for high predation risk, and these yield two very different evolutionary equations because the first involves only a simple energetic equation whereas the second explicitly involves a cognitive demand as well as an energetic demand. The structure of the alternative hypotheses have the form:

1. Predation risk selects for larger body size [and brain size simply readjusts later]
2. Predation risk selects for larger group size, and group size in turn selects for larger brain size (because of the cognitive demands in coordinating large groups [25,59]).

These may both be alternatives (some taxa may prefer one strategy over the other, especially if one is less costly than the other), or they may occur sequentially in a lineage if, for example, increases in body size as a result of the first create spare energetic capacity that enables the second. To test these possibilities, I compare contrast residuals to a well established index of predation risk (degree of terrestriality) and to an index of dietary efficiency (percentage of fruit in the diet).

Group size data for individual species are sourced from [73]. Diet data are sourced from [74], subject to a correction for *Macaca* given by [75]. For the reasons given by [25,28,76–77], actual predation rates are inappropriate for testing hypotheses about the historical impact of predation on the evolution of counter-strategies. Actual predation rates represent the unresolved predation risk that the animals have been unable to control by their evolved anti-predator strategies; it is the intrinsic environmental predation *risk* that drives the evolution of anti-predator strategies. Predation risk is a product of the local predator density, the availability of trees as refuges and body size [77–78]. Since body mass forms part of the brain/body mass residuals, I use an index of terrestriality as a reliable proxy for predation risk (terrestriality exposes primates to higher predation risk with fewer refuges [79]). Contrasts in terrestriality were defined as same or different in terms of Heldstab et al.’s [80] 3-point categorical classification, with further reference to the primary literature for more nuanced interpretation in cases where both species fell into the same category.

To determine whether the data form natural clusters (statistical grades, as opposed to taxonomic grades), I used *k*-means cluster analysis. Inspection of the data suggests that within-genus contrasts form a distinct cluster on the left of the graph, whereas contrasts formed by comparisons of species belonging to different genera (in many cases, different families and even sub-orders, which inevitably have deeper divergence dates) form a scatter of datapoints to the right of the graph. To avoid confounding grade with taxonomic effects, I used as the baseline from which to calculate residuals for the cluster analysis the RMA regression for within-genus contrasts only. This sets the regression up through the centre of the data distribution, rather than across it as is inevitably the case when OLS regression is used with data that have high error variance [25,28]. It also focusses the analysis of residuals on the initial post-divergence phase, thereby avoiding any quadratic effects that might arise later through convergence back onto the allometric regression line.

Cluster analysis makes no assumptions about the form of the data; instead, it simply seeks to identify natural breaks within the distribution of the data. There are a number of clustering methods available (DBscan, Jenks, Gaussian mixture models, head/tail breaks, *k*-means, Clauset-Newman maximum likelihood method, broken stick models) and we have used all of these with different kinds of biological data [25,28,81–85]. Where different methods have been run on the same data, they produce very similar results. Here, I use *k*-means cluster analysis, mainly for practical convenience and because it is conceptually simpler.

There are no formal tests for finding *k* (the optimal number of clusters). Goodness of fit inevitably increases with *k* and, by definition, eventually approaches r^2^=1.0 when *k* is equal to the number of data points. Convention is to find the value of *k* that maximises fit while minimising the number of clusters, subject to the rule that there should be as few clusters as possible consisting of a single datapoint. To determine the optimal number of clusters, we can plot the *F*-statistic as an index of goodness of fit against cluster number. Since this will usually be sigmoid or asymptotic in shape, the point of statistical independence can be identified as the point at which the slope starts to decrease (the point conventionally identified as the ‘break’ in slope in a ‘broken stick’ model). A more sophisticated approach notes that, on any asymptotic curve, the point of inflection (i.e. the point where the magnitude of the change in Y starts to decline relative to the change in X) is defined by the point on the X-axis that corresponds to the point on the Y-axis that is 1/e^th^ down from the asymptotic value [86]. I use the latter criterion since it is mathematically more precise and is easier to calculate; it yields results that are identical to the broken stick method [28].

One central concern is the fact that the way OLS regressions are calculated causes estimates of the slope to tend to b=0 if the assumptions of the OLS regression model are violated (i.e. when the data are not bivariate normal, there is significant error variance in the estimates for the X-axis values and/or r^2^<0.95 due to grades in the data causing high error variance on the Y-axis. To determine whether or not Deaner & Nunn’s negative result is an artefact of using OLS regression methods, I ran both OLS and RMA regressions on the data. Although there are still no formal methods for testing whether the slope and intercept parameters of an RMA regression equation differ from the null hypothesis of a=b=0, we can do so indirectly by a two-step process in which we ask first whether the OLS parameters differ from a=b=0 and then whether the RMA parameters differ significantly from the OLS parameters, using the standard errors on the OLS intercept and slope to calculate the t-statistic in the usual way (i.e. we treat the OLS parameters as the null hypothesis). When r^2^≥0.95, RMA and OLS regression methods converge and give the same answer, and the extent to which the two regression methods agree therefore provide *prima facie* evidence for grades in the data.

Where a directional prediction is being tested, 1-tailed p-values are given and so indicated; all other statistical tests are 2-tailed. Where separate tests are run on different grades, I use Fisher’s meta-analysis for small samples [87] to combine the results: this is a maximum likelihood test that uses 1-tailed p-values to ask how likely it is that the set of test results would be as extreme as those observed if there were no underlying trend in the data (with df = twice the number of tests sampled).

The data are provided in online Tables S1 (for primates) and S2 (for hominins).

## Results

The analysis proceeds in three steps. First, we seek to ascertain whether or not there is a brain lag effect, given the availability of more accurate divergence dates and the use of more appropriate statistical methods. There are three possible outcomes: (i) zero correlation if Deaner & Nunn were right and there is no lag effect (i.e. brain and body size always change together in a tight co-evolutionary ratchet, as has been shown to be the case for the group/brain size relationship in primates [18]; (ii) a significant positive correlation if there is a lag effect in which body size changes first (as predicted by [1]); and (iii) a significant *negative* correlation if there is a lag effect, but it is brain size that changes first (in effect, a body size lag). If the answer is (ii) or (iii), we proceed to a second step and ask whether the regression line stably approaches the allometric line (i.e. a simple lag effect) or extends beyond it (i.e. something else is going on). If the answer is the second, then, as a third step, we ask (i) how does the species concerned solve the energetic shortfall that is an inevitable consequence of increasing brain size and (ii) why does this pattern occur rather than a simple lag effect.

### Is there a brain lag effect?

I first consider the original Deaner-Nunn analysis with the Purvis divergence dates to check whether an RMA regression changes their results. If the data are bivariate normal and r^2^>0.95, then an RMA regression will produce exactly the same best fit equation as an OLS regression. If, on the other hand, there are grades in the data (and r^2^<0.95 as a result), then RMAτOLS. In this case, the RMA regression will always give a truer estimate of the slope. I then ask whether more recent dating of divergences makes a difference.

The original Deaner-Nunn data with the Purvis divergence dates are plotted in Fig. S1. As Deaner & Nunn found, the regression is positive but does not differ significantly from *b*=0 (b=0.002±0.004sem, standardised *β*=0.101, r^2^=0.010, F_1,23_=0.23, p=0.316 1-tailed). However, an RMA regression of these data (Residual = -0.166 + 0.018*Date) yields a significantly steeper slope than the OLS regression (t_23_[0.018=0.002±0.005sem]=41.0, p<<<0.0001), with a significantly more negative intercept (t_23_[-0.166=-0.007±0.051sem]=3.12, p=0.0024). In other words, the regression model used makes a significant difference. One likely reason is that the dataset is not homogenous, but consists of a set of grades.

Fig. 1 plots the residuals for the same contrasts in brain mass regressed on body mass against the updated Perelman divergence times. The solid horizontal line gives the null hypothesis (slope b=0), and the heavy hatched line is the OLS regression. The OLS regression with the Perelman divergence dates is significant (b=0.005±0.002sem; standardised *β*=0.445; r^2^=0.198, F_1,23_[*β*=0.445>0]=5.68, p=0.013 1-tailed). Note that the intercept is negative (body size changes first), though not significantly so (*a*=-0.057±0.045sem; t_23_=-0.1.26, p=0.111 1-tailed). The RMA regression through these data is: Residual = -0.1574 + 0.0113*Date. Its slope is significantly steeper than that of the OLS regression (t_23_[0.0113=0.005]=3.15, p=0.0022 1-tailed), and the intercept significantly lower (t_23_[-0.157=0.057]=2.23, p=0.018 1-tailed; and hence also significantly below 0). In this case, the RMA and OLS regressions are both significant: more accurate dating of divergence times significantly improves the outcome estimates. As before, an RMA regression yields a significantly steeper slope with a more negative intercept than an OLS regression, again suggesting that there may be grades in the data.

**Fig. 1.**
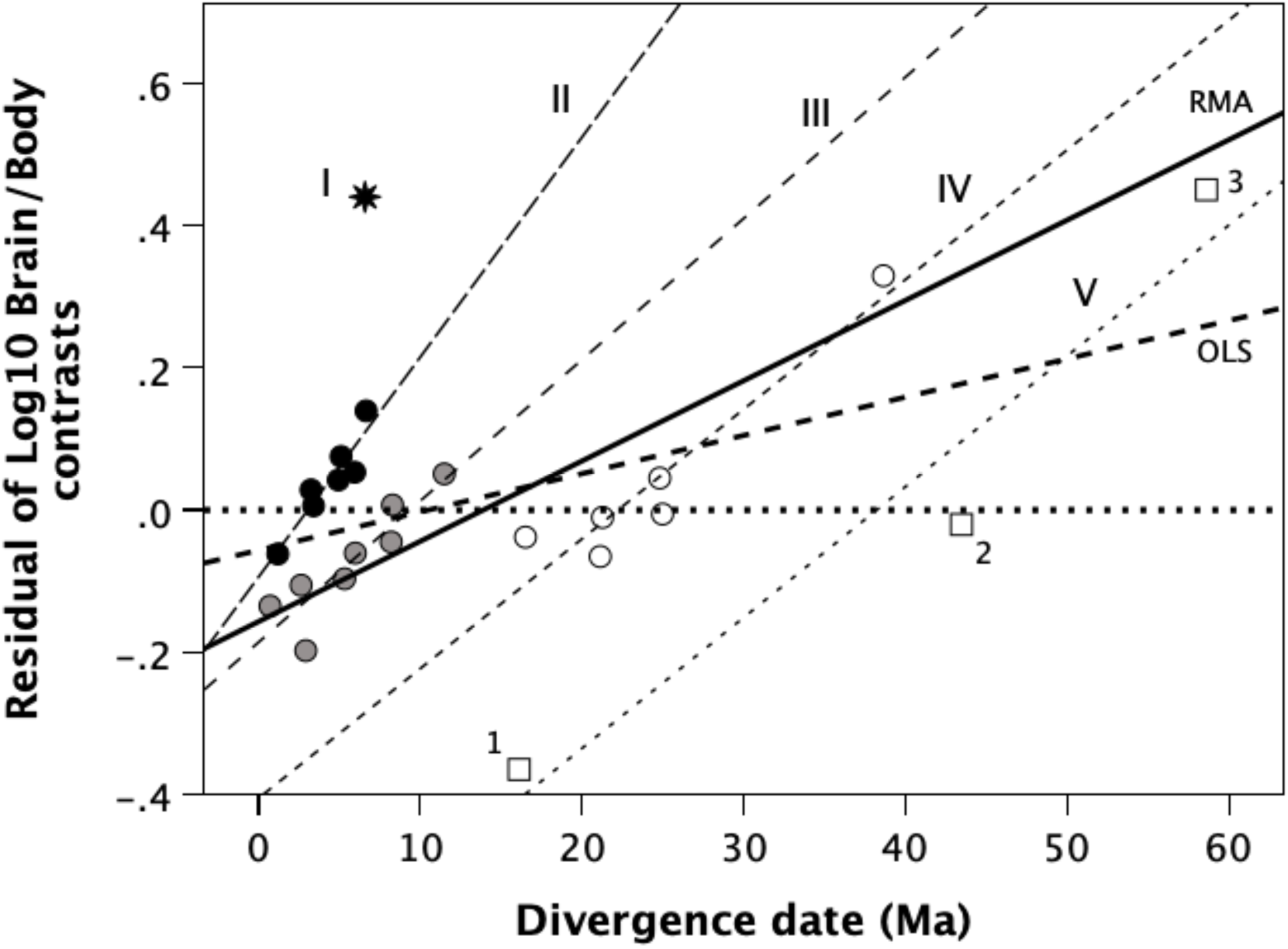
Residuals of the contrasts in log_10_ brain mass regressed on contrasts in log_10_ body mass, plotted against time of divergence for the species pairs (based on the Perelman phylogeny). The horizontal dotted line is the null hypothesis (b=0); the heavy dashed line is the OLS regression for the plotted data; the heavy solid line is the RMA regression. A *k-*means cluster analysis of residuals from the within-species RMA regression line indicates that the data form five distinct grades; the contrasts allocated to the grades by the clustering algorithm are indicated by the different symbols (and the Roman numerals by the respective grade OLS regression lines). Star: grade 1 (*Homo-Pan*); filled circles: grade 2 (within-genus contrasts); grey circles: grade 3 (within-genus contrasts); unfilled circles: grade 4 (between-genus contrasts); squares: grade 5 (between-family contrasts) (1: *Aloutta/Lagothrix*; 2: *Piliocolobus/Leontopithecus*<u>; 3: *Daubentonia/Avahi*)</u>.

To determine whether the data are better described as a series of statistical grades, I ran a *k*-means cluster analysis on the data from Fig. 1, using the RMA regression for the within-genus contrasts as the baseline against which to calculate residuals: Residual [within-genus brain/body] = -0.14106 + 0.017439*Date [1] With residuals calculated from eq. [1], a *k-*means cluster analysis was run for 2≤*k*≤7. All cluster divisions tested provide significant fits to the data (ANOVA, p<0.001). Our problem is a model-fitting one: how to find the clustering pattern that maximises fit while minimising the number of clusters. For this we find the phase transition point in the distribution of goodness of fit as a function of *k* (number of clusters). Fig. S2 shows that this is *k*=5.

Fig. 1 plots the five clusters identified by this method as different symbols (and the associated Roman numerals). Since r^2^≈0.95 for individual grades, OLS regressions are shown in each case. One cluster (the *Homo-Pan* contrast) contains just a single member. Pairwise comparisons between successive grades indicate that, as a set, the slopes parameters do not differ significantly across the four grades (Fisher’s test, p=0.089), but the intercept parameters do (p<0.0001) (Table 1). Notice that all but two of the datapoints in the two leftmost grades (labelled II and III) are congeneric species; and all but one of the datapoints in the two rightmost grades (labelled IV and V) are comparisons between species that belong to different genera, families or suborders.

**Table 1.**
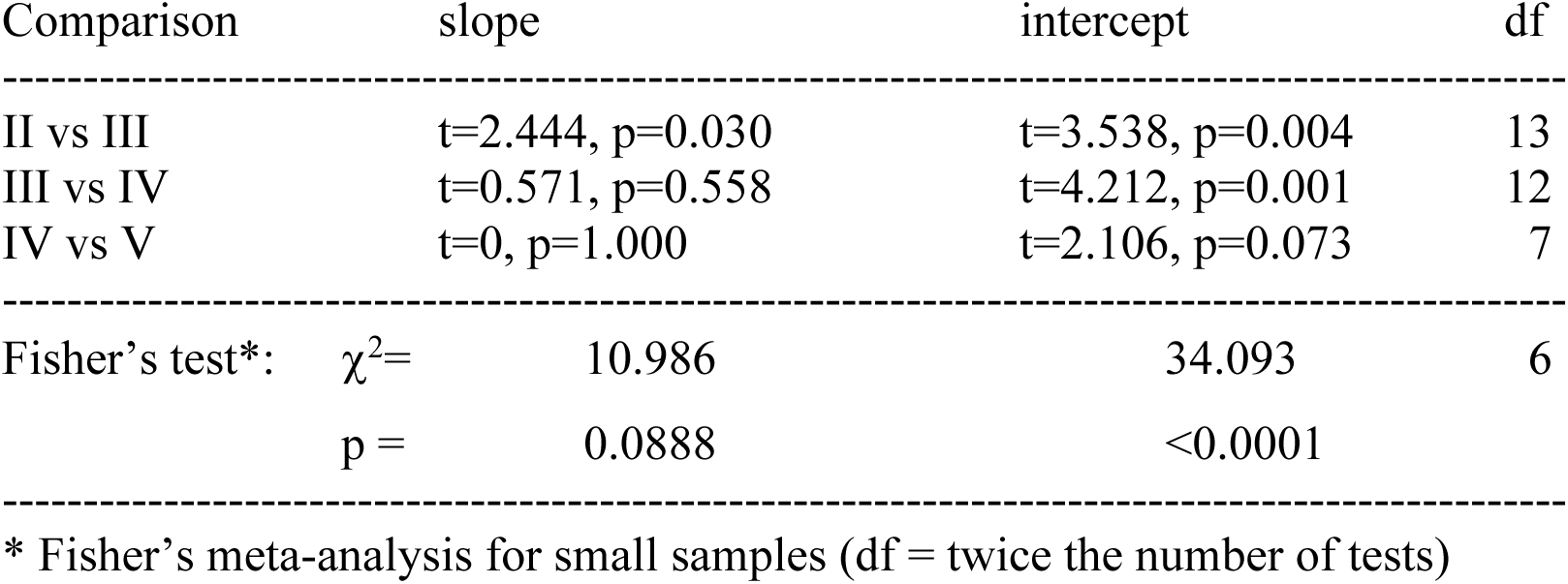
Pairwise comparisons of slope and intercept coefficients of the regression equations for the grades in Fig. 1.

Grades II-V have individually positive and significant or near-significant regression slopes. The mean correlation across the four grades is r=0.933 (taking grade into account, overall r^2^=0.892, mean standardised β=0.936: Fisher’s procedure for combining independent tests: χ^2^=19.2, df=2*4=8, p=0.013, indicating a consistent underlying positive trend). As is invariably the case when grades are present, failing to take them into account results in an underestimate of the true regression slope, as well as the goodness of fit [25,28].

Three points may be noted about the distributions in Fig. 1.

First, in each grade, the more recent contrasts have a negative residual, while the older ones have positive values. For the two mainly within-genus grades (II and III), the distribution of datapoints in grade II is displaced upwards compared to the earlier grade III: in grade II, only 1/7 datapoints falls below the zero line (where brain size is what would be expected for body size), but 6/8 do so in grade III. In other words, the intercepts become increasingly more negative as the divergence time of the grade gets deeper (i.e. more rightwards on the graph) (Fig. 2a; r^2^=0.999, standardised β=-1.0, t_2_=-47.9, p=0.0004 2-tailed). The regression equation for the data in Fig. 2a is significantly different from the null hypothesis of no correlation (i.e. slope b[H_0_]=0) (r^2^=0.999, standardised β=-1.0, t_2_=-47.9, p=0.0004 2-tailed). Notice how tightly the data points cluster along the regression line. This suggests that early divergences favoured increasing body size, whereas later divergences were more likely to opt for increasing brain size (so as to increase group size). In contrast, the slope parameters become increasingly steep in more recently diverged lineages (Fig. 2b: r^2^=0.983, standardised β=-0.991, t_2_=-16.7, p=0.002 2-tailed). The best fit regression is in fact an inverse equation:

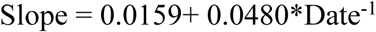

**Fig. 2.**
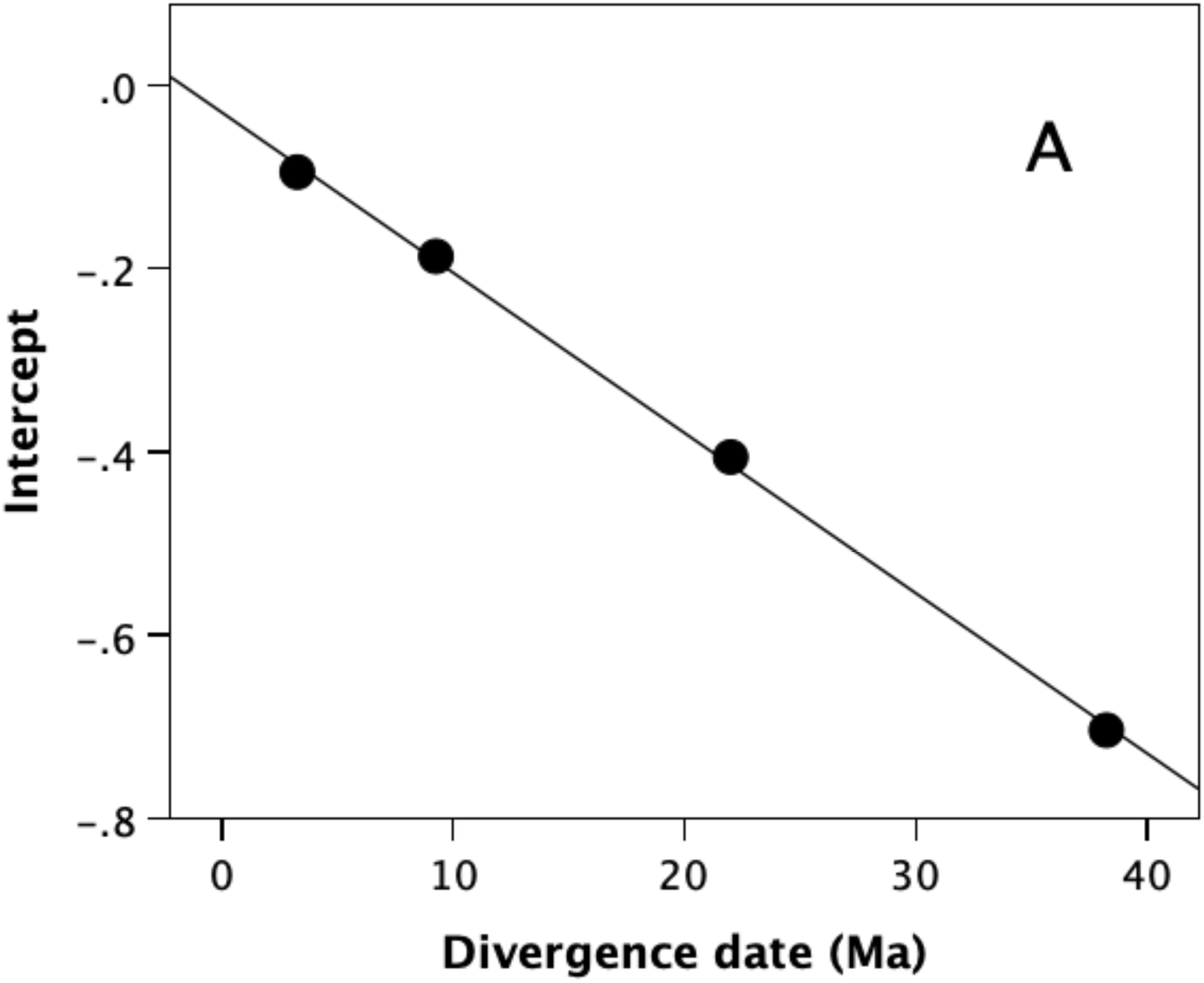
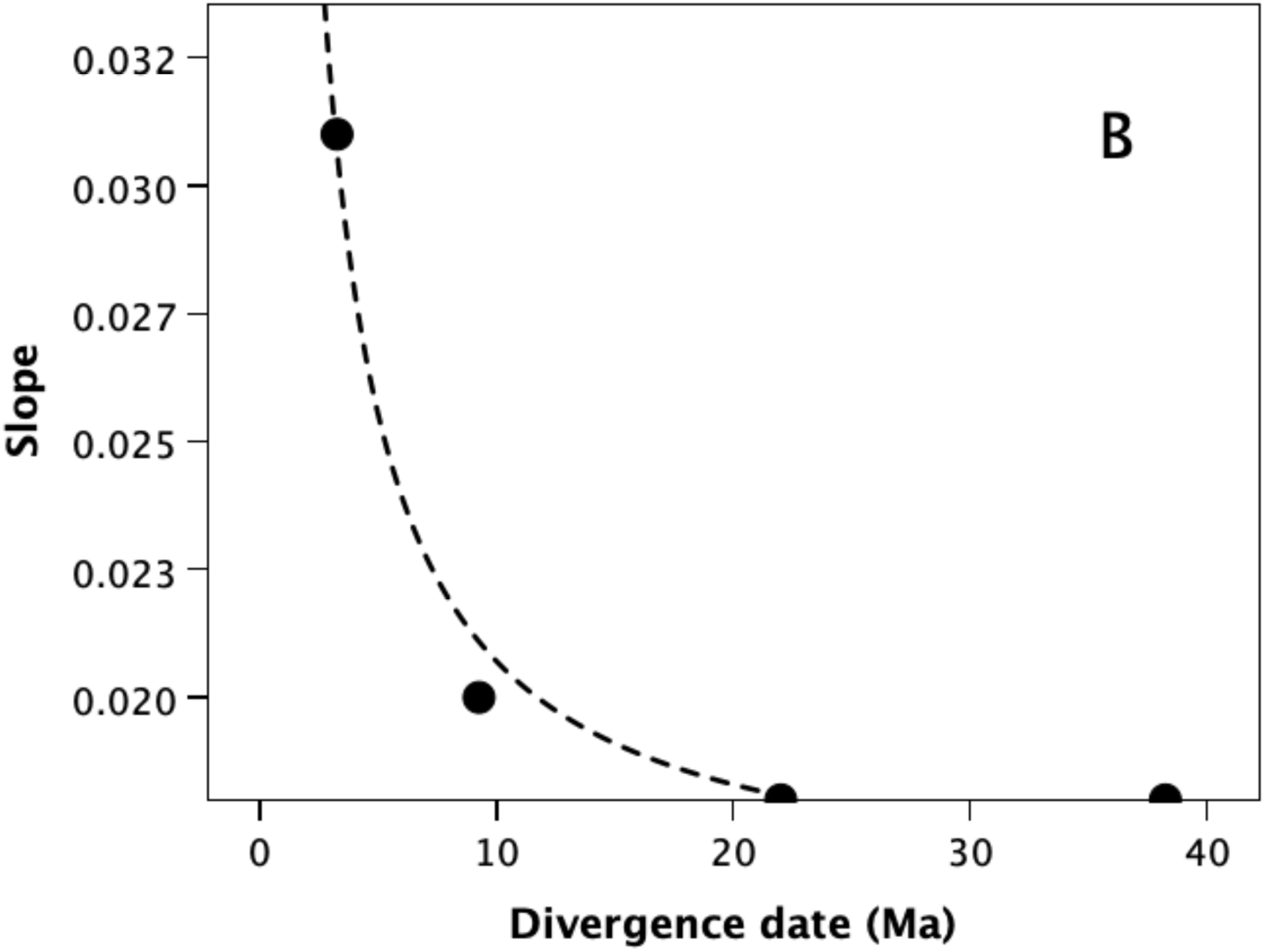
(a) Intercept and (b) slope values for the regression equations for the four grades in Fig. 1, plotted against mean Perelman divergence dates (indexed by the date at which the grade’s regression line crosses the origin line at ratio = 0). The mean divergence date is calculated as the date at which the grade OLS regression crosses the Y=0 line.

(r^2^=0.983, t_2_=10.67, p=0.002 2-tailed). This suggests that younger diveregences favoured investing in larger brains rather than larger bodies. One explanation for this might be that more recent grades are able to exploit the energetic savings generated by their ancestors’ earlier increases in body mass. In effect, increases in body mass have provided a platform off which it has been possible to divert energy into brain growth, thereby catapulting the species concerned onto a higher cognitive plane.

The second point to note is that the time lapse to converge on the allometric line is 2.0 million years for grade II and 8.6 million years for grade III. Notice, also, the high value for the *Homo/Pan* contrast (which Deaner & Nunn chose to discount from their analysis, possibly because it was so divergent). Its position suggests that it might represent the end-point of a separate grade whose starting point is much higher than that for grade II (see below).

The third point is that, contrary to the original brain lag hypothesis, the brain/body ratios for all the grades does not stabilise back on the regression but instead continues to diverge past it in favour of increasingly large brains. The contrasts for grade IV are phylogenetically much deeper. All but two of the contrasts in this grade are strepsirrhines (the other two being New World monkeys and a great ape pair, both with unusually deep divergence times). Since the strepsirrhine contrasts cluster around the zero line, they appear to involve a straight forward brain lag that has converged on the allometric regression. It seems only to be anthropoids that extend beyond the regression line into positive territory (grades I-III). This is in line with evidence that the strepsirrhines do not exhibit a strong social brain relationship [19,24–25,53].

### How the energetic constraint was resolved

A significant deviation from the common allometric regression line necessarily creates an energetic demand to fuel the additional body or brain growth. Of these, brain growth is the more important because brain tissue is ∼20 times more energetically costly than somatic tissue (the expensive tissue hypothesis [88–91]. One solution to this is to switch to a more nutrient-rich diet. Primates broadly divide into folivores and frugivores, with frugivory considered the richer, more energy-accessible diet (mainly because a folivorous diet requires fermentation by specialised bacteria to enable the animal’s digestive system to access the nutrients contained within the cell walls of leaves [92].

To test whether the switch from investing in body size to investing in brain size is associated with a change in diet, Fig. 3 plots contrasts in percent of fruit in the diet against the residual of contrasts in brain size regressed on contrasts in body size. The OLS regression is positive and significant (r=0.415; F_1,22_=4.59, p=0.022 1-tailed): species with larger brains for body size are more likely to be frugivorous. I then compared the percentage of fruit in the diet separately for contrasts where brain size residuals were positive versus negative (Fig. S3). Overall, the difference is not significant (t_22_=1.06, p=0.151 1-tailed). However, most strepsirrhines are insectivorous and belong to a smaller-brained sub-order than haplorrhines. Excluding strepsirrhines yields a significant difference within the anthropoids (t_16_=2.05, p=0.029 1-tailed). This suggests that when the anthropoids opted for an increase in brain size, a change in diet to a more nutrient-rich regime was required, but this was not the case when they opted for an increase in body size – presumably because the energy demand was much lower. It seems that a dietary adjustment of this kind was not necessary for the strepsirrhines, perhaps because their much smaller brains require less energy and because they do not deviate so far from the allometric regression (in other words, after increasing body size, they simply converge back onto the allometric regression line).

**Fig. 3.**
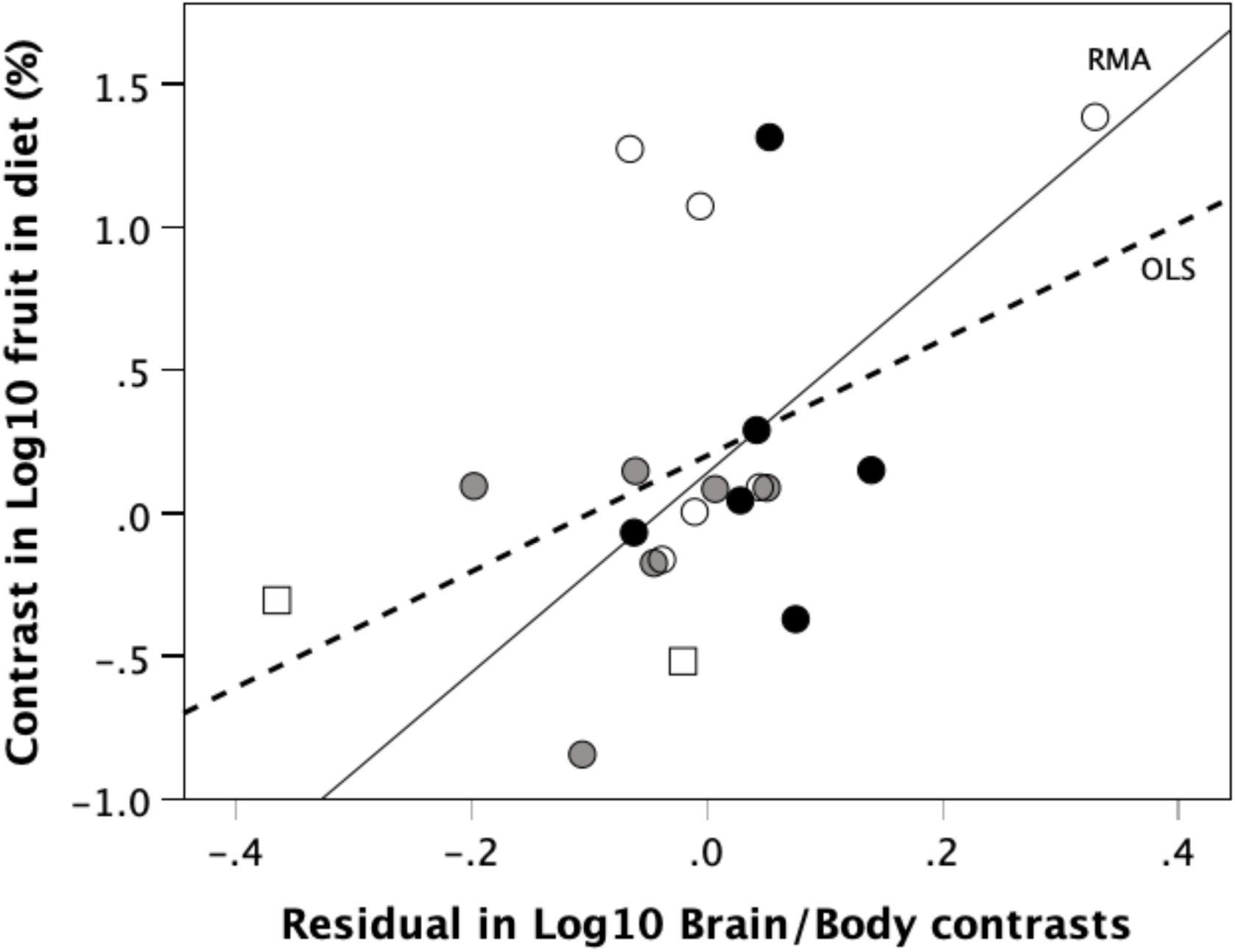
Contrast in percent fruit in diet plotted against residual of contrast in log_10_ brain volume from contrast in log_10_ body mass. Dashed line (OLS): ordinary least squares regression; solid line (RMA): reduced major axis regression. Symbols as for Fig. 1.

### Was predation risk the selection pressure involved?

To test the hypothesis that the underlying driver for the changes in brain and body size is predation pressure, I plotted contrasts in brain:body mass as a function of contrasts in the terrestriality index (Fig. 4). For contrasts where there is no difference in terrestriality, residual brain size values cluster around 0 (brain size residuals do not differ), but residual brain size is significantly different where one of the pair is more terrestrial (t_21_=2.61, p=0.016). This suggests that predation risk is likely to have played a role in triggering changes in relative brain or body size.

**Fig. 4.**
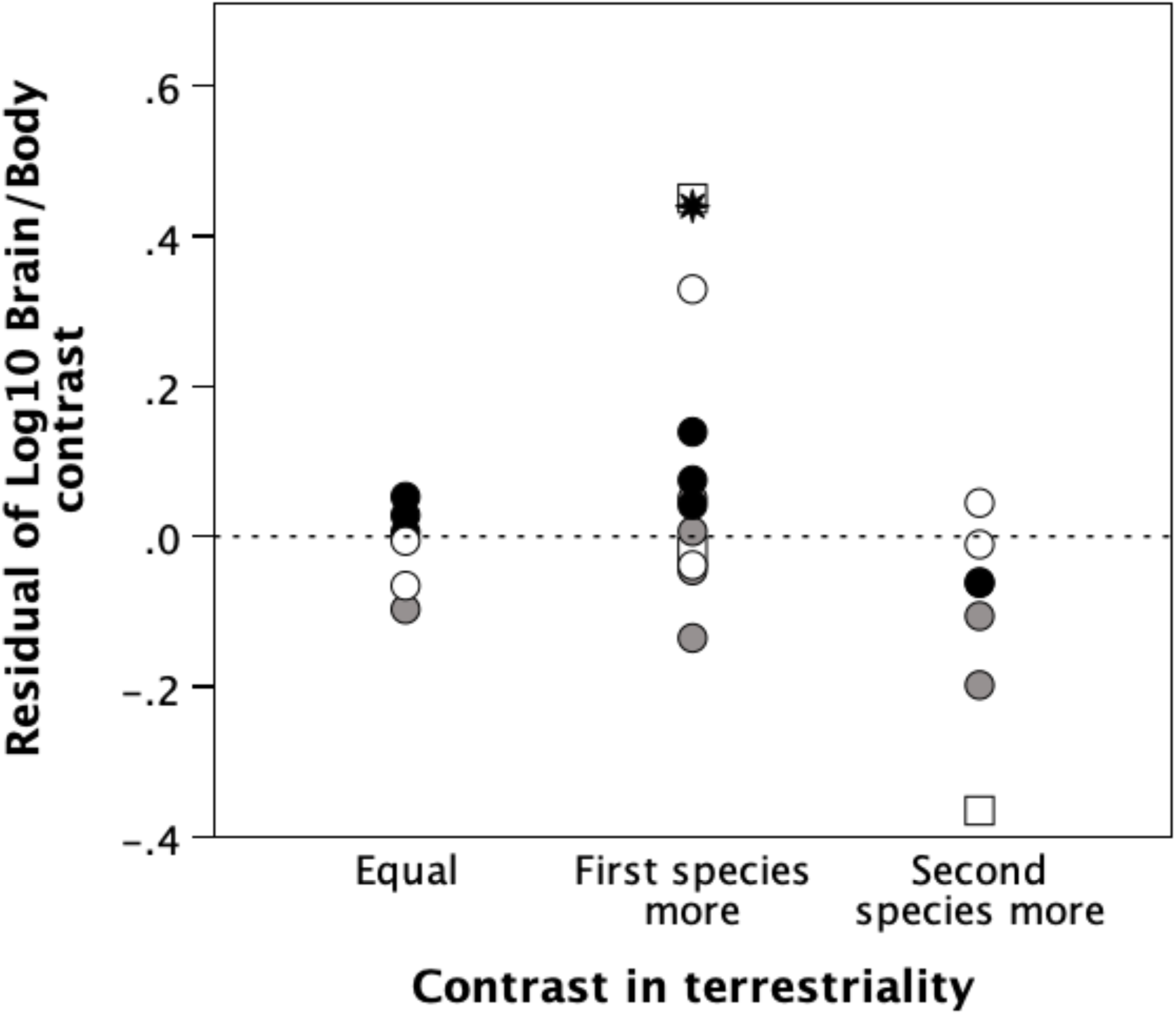
Residuals in the contrasts in log_10_ brain size regressed on log_10_ body size as a function of the relative degree of terrestriality. Dashed horizontal line indicates residual = 0 (brain size is directly proportional to body size). Symbols as for Fig. 1.

As a defence against predators, primates typically adopt either large body size (e.g. gorillas, hominins) or large group sizes (most other diurnal species). Deaner & Nunn [1] tested for a relationship between social group size and residual brain size, but found none. However, using more up-to-date data on species mean group sizes from [73], there is in fact a highly significant positive overall correlation (Fig. 5: r=0.612, p=0.005 2-tailed). All three grades where N>3 exhibit positive slopes (r=0.806, p=0.015; r=0.410, p=0.312; r=0.724, p=0.052; Fisher’s procedure: χ^2^=16.64, df=6, p=0.011, indicating a consistent positive trend across the three subsamples).

**Fig. 5.**
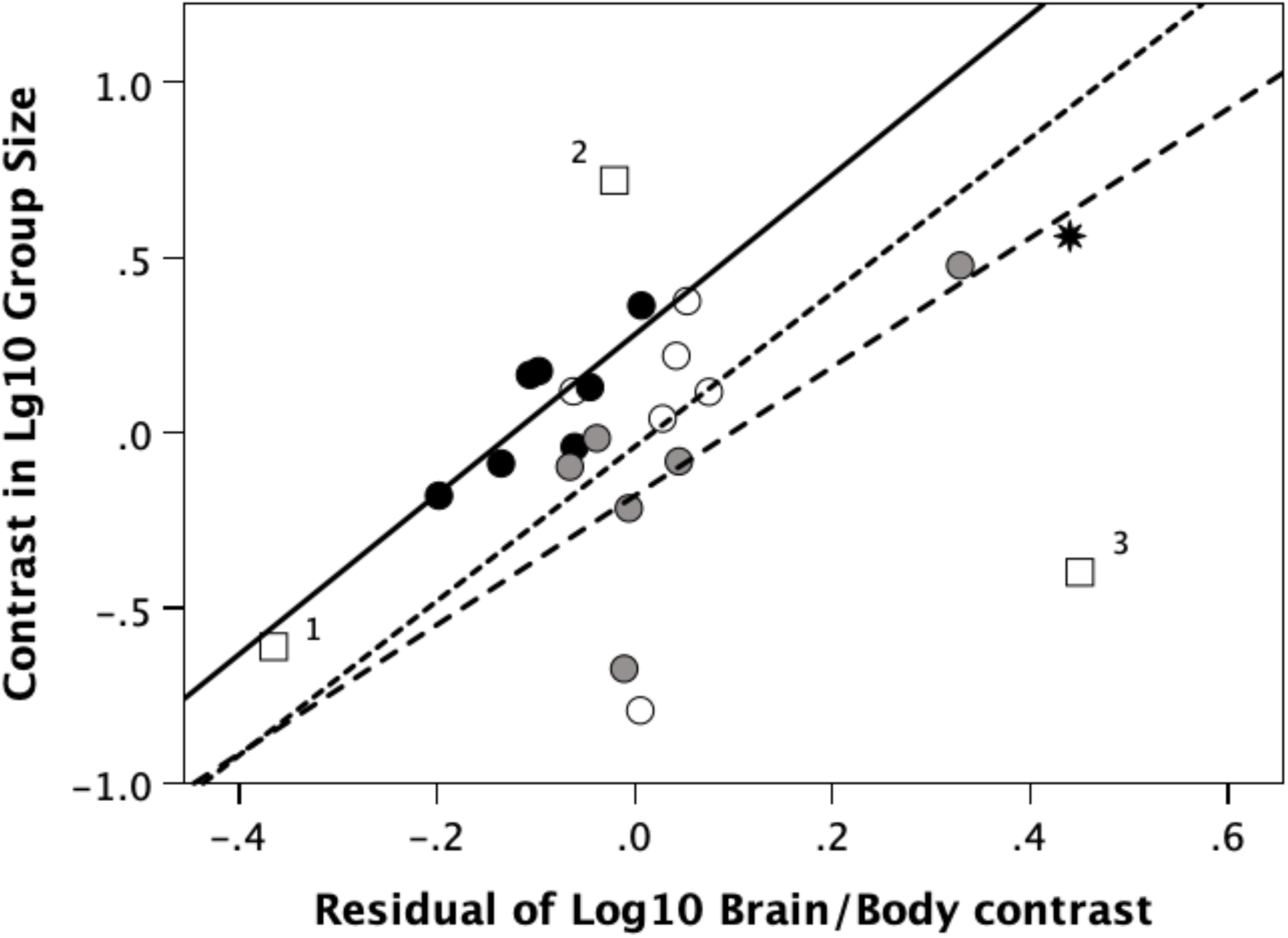
Contrasts in log_10_ mean group size plotted against residual of contrast in log_10_ brain mass regressed on contrast in log_10_ body mass. Symbols as for Fig. 1. Regression lines are for grades II (solid line), III (dashed line) and IV (dotted line), as defined in Fig. 1.

Note that two of the squares (grade V) in Fig. 5 lie well off-diagonal, which is usually indicative of a grade shift. The higher one (datapoint 2) represents a contrast between *Piliocolobus* (a semi-terrestrial Old World monkey) and *Leontopithecus* (an arboreal New World monkey) and indicates a shift between an already known highly social and a much less social grade [25] as well as being a very ancient phylogenetic separation. The lower datapoint (3) is a contrast between two distantly related strepirrhines (*Daubentonia* and *Avahi*). For present purposes, and to be conservative, I followed Deaner & Nunn [1] and others in assigning a group size of N=1 to *Daubentonia* on the grounds that it usually forages alone. However, there are strong grounds for believing that this species actually lives in dispersed communities of N≈8 [28]. If their correct group size is anywhere close to 8, this would place this contrast very close to the star (*Homo-Pan* contrast), yielding an slightly improved overall linear fit (r^2^=0.622, p=0.004) – the important point here being that even if this datapoint is subject to a great deal of measurement error it doesn’t destabilise the overall relationship.

Deaner & Nunn [1] implicitly assumed a causal relationship in which changes in body size drive changes in brain size, with changes in group size being presumably at best a default by-product of changes in brain size. A path analysis of the relationships between these three variables (based on significant standardised slopes in multiple regressions with each variable in turn as the dependent) yields a best fit model in which brain size is correlated independently with body size and group size, with no relationship between body size and group size (Fig. 6). Importantly, mediation analysis for the six possible three-way causal pathways indicates that the only significant pathway is group size predicting brain size, and brain size predicting body size (Sobel test: p=0.028); none of the other alternative pathways are significant (p≥0.103). In other words, group size selects for an increase in brain size, and a larger brain then selects for an increase in body size so as to exploit Kleiber’s Law to reduce the energetic costs. This would seem to suggest that, aside from the exceptions noted above, body size is only rarely exploited as a first line defence against predators, at least by anthropoid primates. Rather, most species opt directly for group-living as the solution, with body size being one important way in which the energetic demands of brain growth can be offset.

**Fig. 6.**
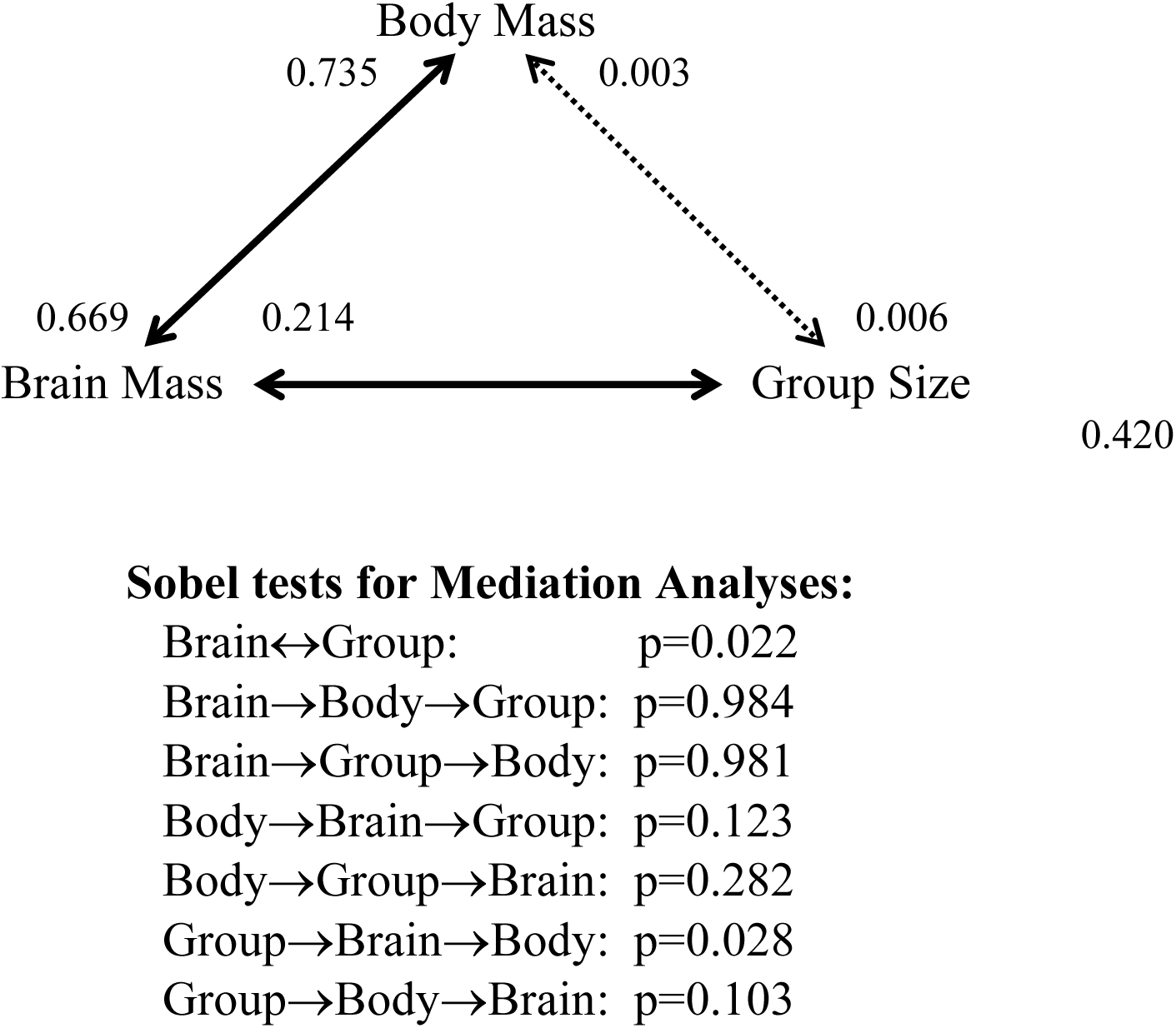
(a) Path analysis of the functional relationships between contrasts in log_10_ brain mass, contrasts in log_10_ body mass and contrasts in log_10_ social group size, based on multiple regressions for each in turn as the dependent variable. The numbers adjacent to the arrow heads are the standardised slopes (ßs) in each direction (where the adjacent arrowhead specifies the dependent variable). Solid lines indicate significant predictors; dotted lines indicate non-significant predictors. (b) Sobel tests of mediation analyses for all six possible causal (i.e. directional) sequences, and for bivariate regression between contrasts in brain size and contrasts in group size.

### The hominins as a test case

The hominin lineage is one of the few cases where the fossil record allows us to trace the evolutionary pathway in considerable detail. *Homo* derives from an ape-like australopithecine root with a body size indistinguishable from that of modern *Pan* [95]. Fig. 7 plots the change over time in the brain:body mass residual from the primate RMA regression line (in this case, for want of any more suitable alternative, using corrected ECV to estimate brain mass). The australopithecines fit comfortably on the grade II regression line (and well within its 95% CIs), but the appearance of *Homo* around 2.5 Ma marks a steep upturn in brain size giving rise to a dramatic increase in the brain:body mass residual. The best fit OLS regression has an inverse form (r^2^=0.724, standardised β=0.851, t_9_=-4.857, p=0.0009 2-tailed). Excluding the two earliest australopithecine datapoints gives a significant linear fit (thin solid line: r^2^=0.978, p<0.0001).

**Fig. 7.**
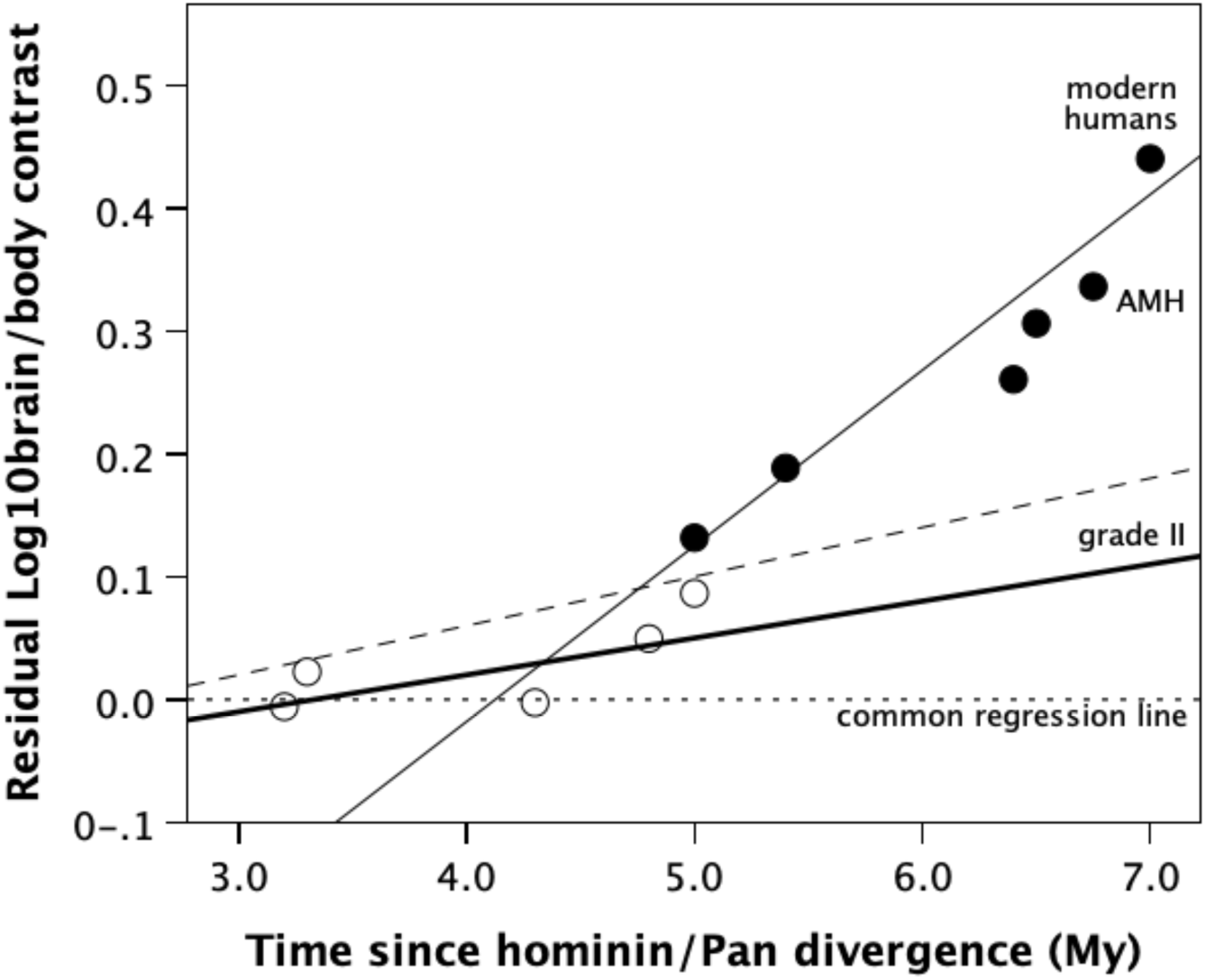
Residual of hominin/*Pan* Log_10_ contrast in brain volume regressed on Log_10_ contrast in body mass for individual fossil hominin species, plotted against time since *Homo/Pan* divergence at ∼7 Ma. The modern human datapoint is the *Homo/Pan* contrast from Fig. 1; the increase in residual brain size compared to fossil *Homo sapiens* (AMH) is due to a marked reduction in human body size in modern populations. Dating is to midpoint of species time span. For this analysis, brain volume is based on ECV, and has been corrected by ∼6% to adjust for the fact that ECV overestimates actual brain mass; in most cases, body mass is estimated from long bone diameter (a reliable index of the body mass an animal has to carry during locomotion). Unfilled symbols: australopithecines; filled symbols: *Homo*. The horizontal dotted line is the common primate regression line. Thick solid line: grade II regression line, with the dashed line giving the upper 95% CI around this regression (from Fig. 1). Thin solid line: OLS regression set to data values >4.0 My. AMH: fossil anatomically modern humans, dating to ∼100 ka.

There is some suggestion that the australopithecines initially undertook a move towards relatively larger body size (residuals below the grade II primate regression line), and then, once the more nomadic early *Homo* began to occupy increasingly open (high risk) environments from around 2.5 Ma, hominins underwent an explosive increase in brain size without corresponding changes in body size. This correlates closely with a dietary shift towards a more nutrient rich meat-based diet from ∼2.0 Ma, initially as scavenging in early *Homo* but later in the form of active hunting among archaic and modern humans from ∼0.6 Ma.

## Discussion

Although Deaner & Nunn [1] found no evidence for a brain lag effect (changes in brain size lagging behind changes in body size) in primates, a statistically more appropriate analysis of their original data, with and without updated divergence times (based on more recent molecular data), suggests that in fact there is a positive correlation as predicted by the brain lag hypothesis. More importantly, an appreciation that the brain data exhibit grade effects rather than being a single homogenous dataset [21–22,24–25] yields a significantly stronger effect. Changes in body size do seem to precede changes in brain size.

However, the subsequent increase in brain size does not involve a simple reconvergence on equilibrium brain/body ratios in the way the brain lag hypothesis assumed would be the case. Instead, at least in anthropoid primates, it involves continued increases in brain size beyond the main allometric regression to a point where brain size is consistently and stably larger than expected for body size. These grades map closely onto the socio-cognitive grades identified by [25]. This result thus confirms the earlier finding that grades within the social brain dataset represent increasing stepwise emphasis on neurally expensive socio-cognitive skills [25,59]. It is important to appreciate that, in both these cases, the grades identified in these relationships are not formally taxonomic in origin in the way that phylogenetic analyses assume. Although there is a modest taxonomic component, they seem to represent attempts by individual species, sometimes genera, to manage the specific environmental conditions they have access to.

These trends in the brain/body relationship seem to be a response to heightened predation risk. Although primates exploit both large body size and large group size as anti-predator strategies [36–40], they face a particular problem in the latter respect: the bonded nature of primate social groups [25,59,93] means that stable changes in group size are difficult to effect because the personalised relationships that underpin these groups involve complex neural pathways [59]. In other words, increases in body size are easier to effect, and hence invariably appear first; once in place, however, they seem to provide an energetic platform that generates sufficient spare capacity derived from savings of scale via Kleiber’s Law to allow species to invest in larger brains. When combined with switches to a more nutrient-dense diet, this allows brain size to break through the brain/body size equilibrium threshold of the allometric regression, permitting much larger groups to form when lineages are able to exploit their ancestors’ investment in increased body size. This trajectory is well illustrated by the historical changes in relative brain size in fossil hominins (Fig. 7).

Two important features of this process are worth emphasising. First, the way the grades in Figs. 1 and S3 are staggered suggests a ratcheted effect in which the earliest transitions largely emphasised body size over brain size, whereas later transitions were able to build on changes already in place to catapult themselves onto a higher cognitive plane in which group size replaces body size as the main anti-predator strategy. Second, the switch from large body size to large brain size appears to take place over a much longer time period than we might expect for most anatomical changes. In the two within-genus grades (II and III), this switch is distributed over periods of 2 and 8 My (Fig. 1). This suggests that, as species seek to extend the range of ecological niches they can occupy into ever more predator-risky habitats, there was a form of directed evolution in response to specific selection pressures which has to act against considerable evolutionary inertia (represented by the costs involved in both the higher energetic demands of larger brains and the need for considerable neural restructuring).

The present results reinforce recent findings by [20–22] that there are shifts in allometric relationship that occur within the mammals (as Jerison [2] originally pointed out]. Across mammals in general, Smaers et al. [21] concluded that these shifts do not always involve cognitive (i.e. brain size) responses to environmental conditions. This is to be expected because the social brain hypothesis is essentially an anthropoid primate effect, with only a small number of non-primate taxa exhibiting it [19,53]. Different mammalian lineages exhibit different slope parameters in the steepness of the brain/body correlation that is best predicted by having bonded social groups [19]. At root, this reflects the fact that different mammalian lineages have adopted radically different kinds of anti-predator strategies. These have included opting for large body size (the great apes and the cetaceans, some ungulates), opting for large social groups (cercopithecine primates, some great apes), exploiting locomotor advantages (many antelope, patas monkeys, gibbons) and crypsis (many small artiodactyls, the nocturnal strepsirrhines), not all of which are necessarily mutually exclusive [36,94]. They do, however, differ in terms of their efficiency as anti-predator strategies and the costs required to evolve them.

These results would seem to be in line with Lande’s [3] finding that the variance in the primate allometric relationship increases with body size, suggesting that the two become progressively decoupled as body size increases [20,34–35]. This might explain how brain size escapes from the energetic constraint imposed by body size in species with large body sizes (mainly those in grades I and II in Fig. 1). What may have been critical in facilitating this was a shift to more nutrient-rich (i.e. frugivorous) diets, itself dependent on being able to cope with longer day journeys since edible fruits are more sparsely distributed than leaves [36,96]. One important implication of this decoupling of brain and body size, especially among the larger-bodied species (as Lande [3] noted), is that, notwithstanding its persistence in many recent analyses, the use of relative brain size (the Encephalisation Quotient) may be seriously misleading [96–99]. It is difficult to know why the concept continues to persist in the literature.

In sum, contrary to Deaner & Nunn’s [1] claim that there is no brain lag effect, there does in fact seem to be strong evidence for the hypothesis. However, at least in primates, this is not due simply to brain size “catching up” with body size. Instead, it involves an initial increase in body size and then what seems to have been a progressively greater increase in brain size, such that it not merely catches up with body size (as the brain lag hypothesis predicts) but in some cases actually extends well beyond the allometric line, placing the taxon onto a new cognitive grade. This is a rather different picture to the classic brain lag hypothesis, of course. Because the cognitive demands of group-living are very substantial [25,59], this switch to exploiting the benefits of group size is only possible if brain size can increase sufficiently to support the cognitive mechanisms involved. The outcome is an explanation that makes more biological sense and offers a more comprehensive explanation not only for the general macro-evolutionary patterns but also for many of the micro-evolutionary patterns within them that are often ignored in comparative analyses.

## Acknowledgment

A preprint version of this article is available on BioRxiv: doi: https://doi.org/10.1101/2024.02.05.578865 [updated 11 September 2024].

## Statement of Ethics

An ethics statement was not required for this study since no human or animal subjects or materials were used. All data are from secondary sources.

## Author Contributions

RD conceived the project, collated and analysed the data, and was solely responsible for writing the paper.

## Data Availability Statement

All data used in this paper can be found in online Supplementary Information Tables S1 and S2. Further enquiries can be directed to the corresponding author.

---

**Figure S1.**
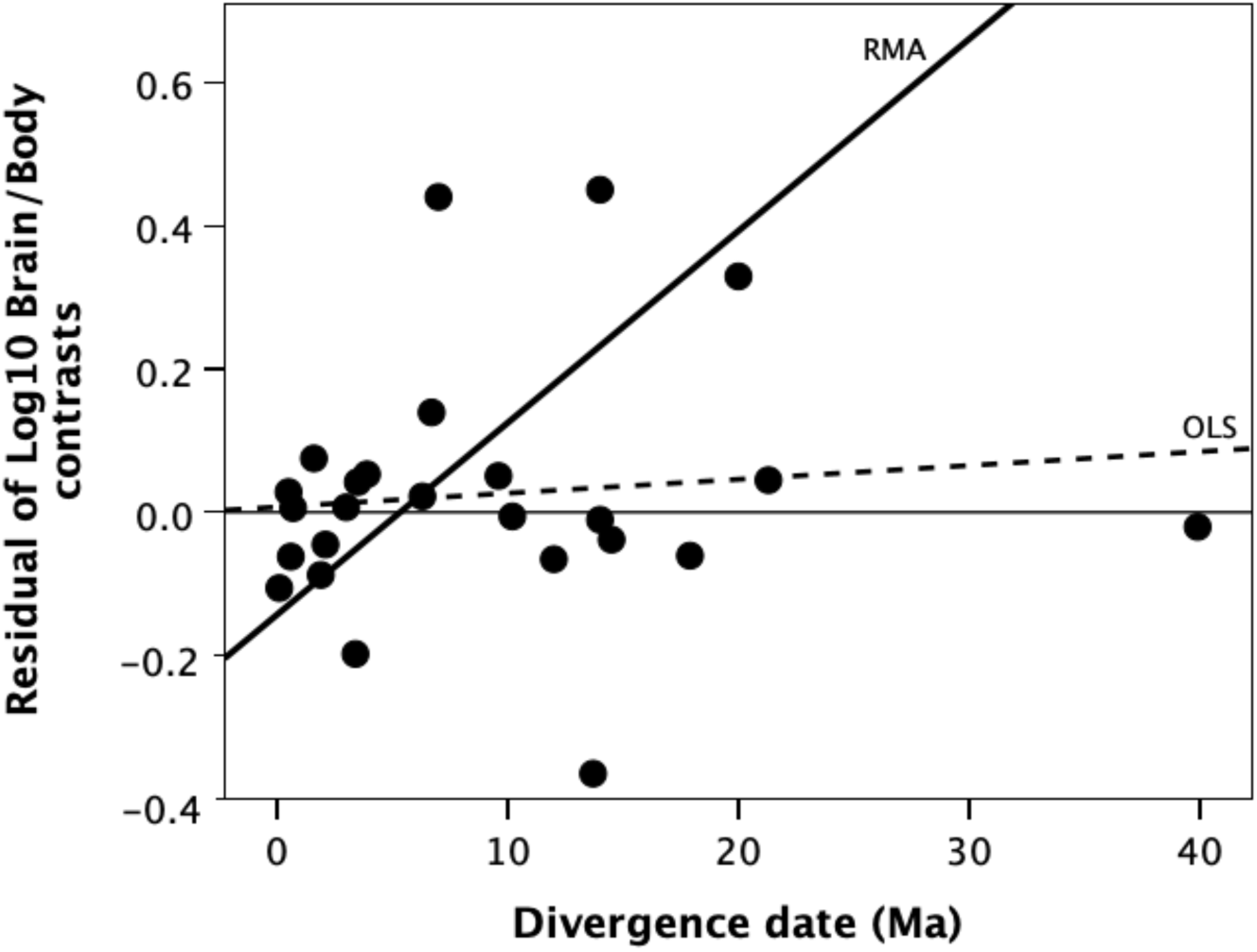
Analysis of contrasts data plotted against divergence time, using the Purvis et al. (1995) phylogeny and estimates of divergence date. The thin horizontal line demarcates a slope of ***b***=0. The thick dashed line is the OLS regression through the data; the thick solid line the RMA regression line..

Figure S1 plots the residuals of the contrast in brain/body mass from the overall regression against the divergence dates from Purvis (1995), as used in the Deaner & Nunn (1995) analysis. The OLS regression through the data give a best-fit regression equation of:

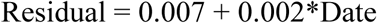

(r^2^=0.010; t_23_=0.488, p=0.631 2-tailed). The regression slope does not differ significantly from 0 (t_23_=0.5, p=0.622 2-tailed). Visual inspection of the data, however, suggests that (aside from the rightmost point) the data suggests a bivariate uniform, rather than a bivariate normal, distribution. The RMA regression has a slope that is significantly steeper than both 0 and the slope of the OLS regression (t_22_=4.467, p≤0.0016), and an intercept significantly below both 0 and the intercept for the OLS regression (t_22_≥-2.337, p≈0.0002). More importantly, the intercept is significantly more negative than 0, as would be predicted by the brain lag effect (i.e. body size changes first). This is strongly suggestive of a dataset that is not bivariate normal, which in turn is *prima facie* evidence for a dataset that has grades.

To determine the optimal number of grades for a *k*-means cluster analysis, we plot a measure of the goodness of fit of the data to different numbers of clusters (in effect, the statistical ‘strain’ in the data) as a function of cluster number across a range of 2≤k≤7 clusters, with goodness of fit indexed as the F-statistic. Since the goodness of fit will at some point inevitably achieve an asymptotic value, we can identify the optimal number of clusters as the point on the X-axis corresponding to the slope transition point in the Y-axis. The latter is identified as the point on the Y-axis corresponding to 1/e^th^ down on the Y-axis from the asymptote.

Figure S2 plots the goodness of fit (indexed as the F-statistic) against number of clusters for the data in Fig. 1 (the plot of residual brain:body mass plotted against Perelman divergence dates). The F-values for all values of *k* are highly significant by a conventional analysis of variance (p<0.001). With an asymptotic value of F≈180, the optimal number of clusters corresponding to 1/e^th^ down from this is 4.5. We round this up to 5.

**Figure S2.**
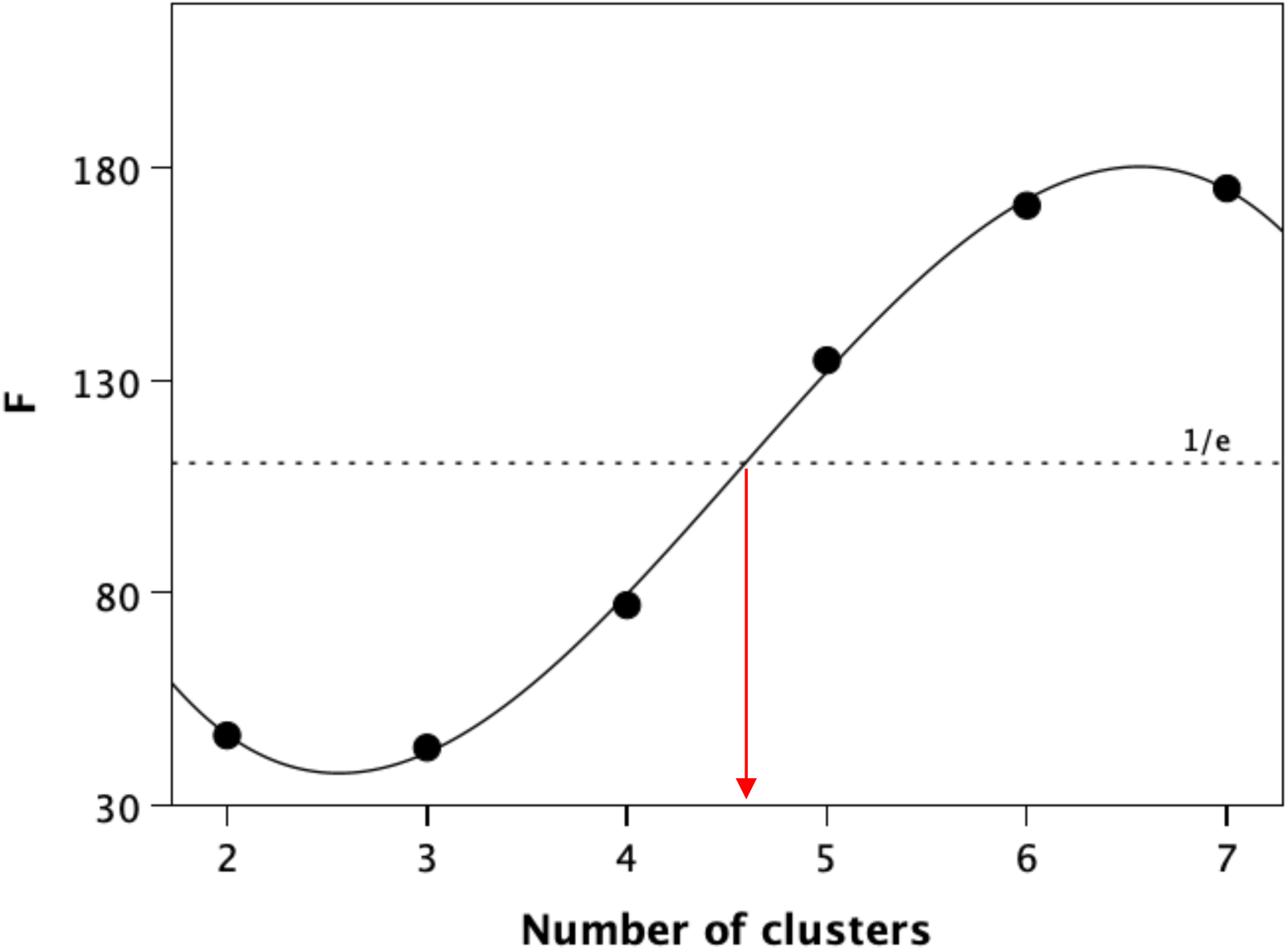
Goodness of fit (indexed by F-value) for number of clusters in a *k*-means cluster analysis. The dotted line demarcates the point of inflexion (1/e^th^ down from the asymptote) that identifies (red vertical arrow) the optimal number of clusters that maximises fit while minimising partitioning of the data.

**Figure S3.**
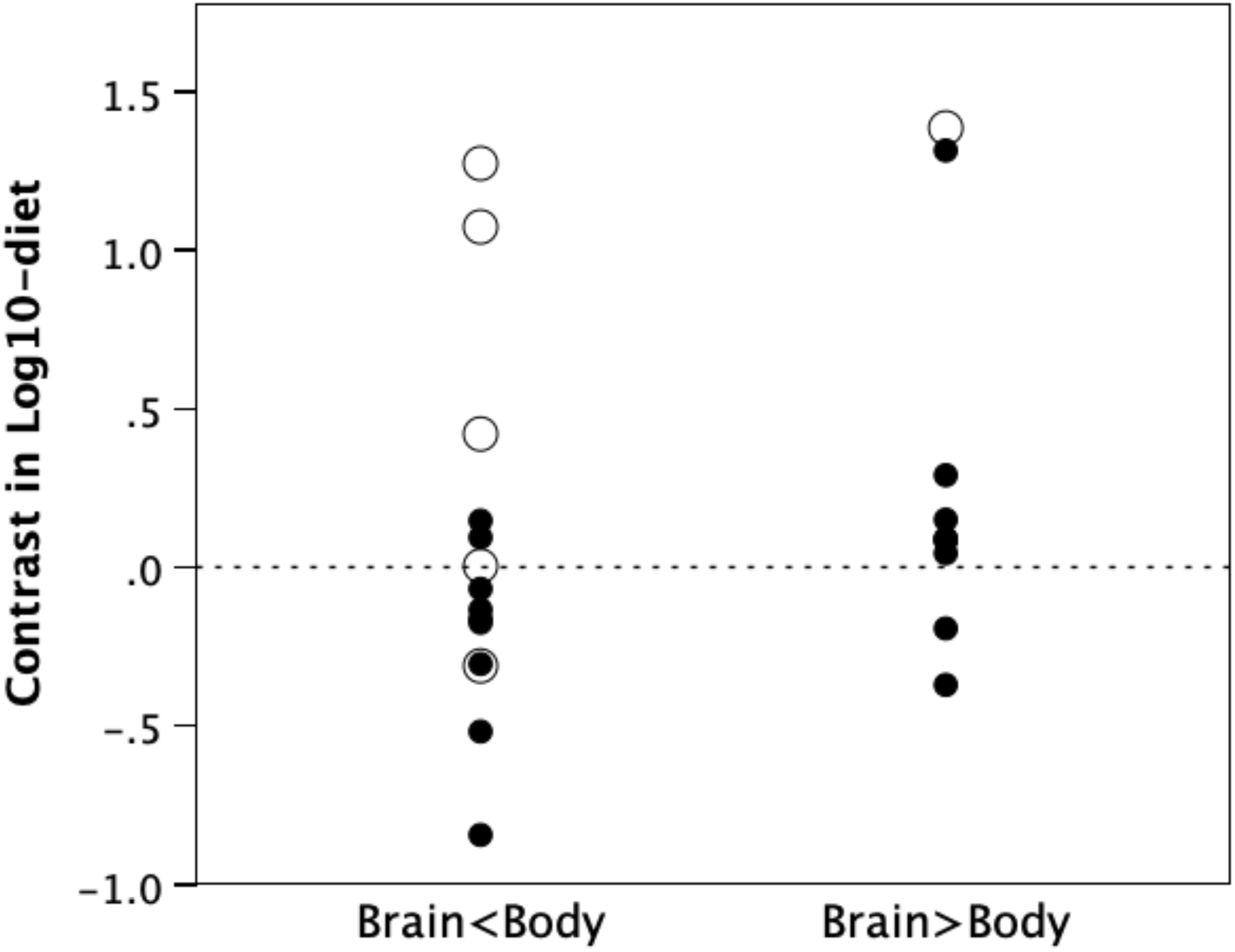
Contrast in log_10_ percentage of fruit in diet for species where residual brain size is negative (brains are smaller than predicted for body size) versus those where it is positive (brains are larger than predicted). Filled symbols are haplorrhines; unfilled symbols are strepsirrhines.

**Table S1.**
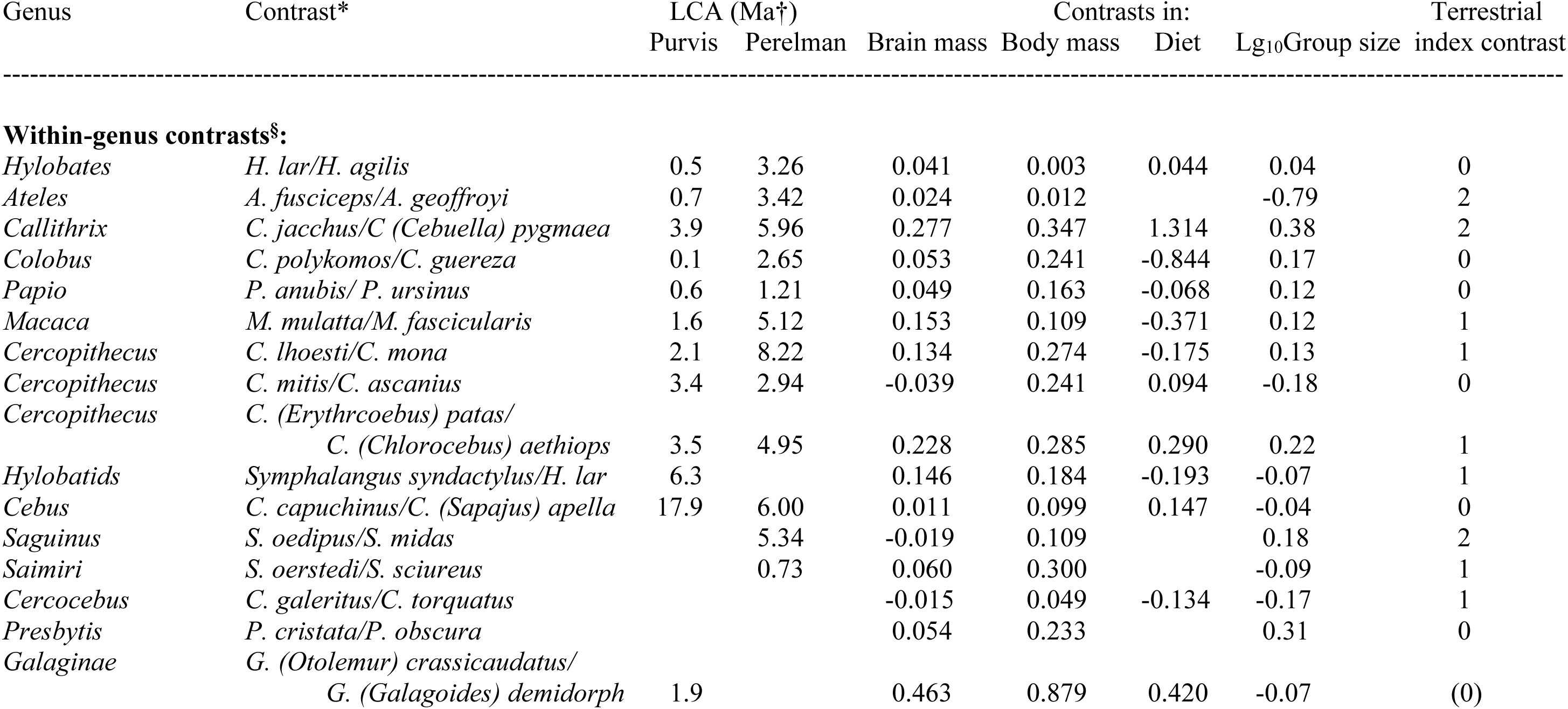
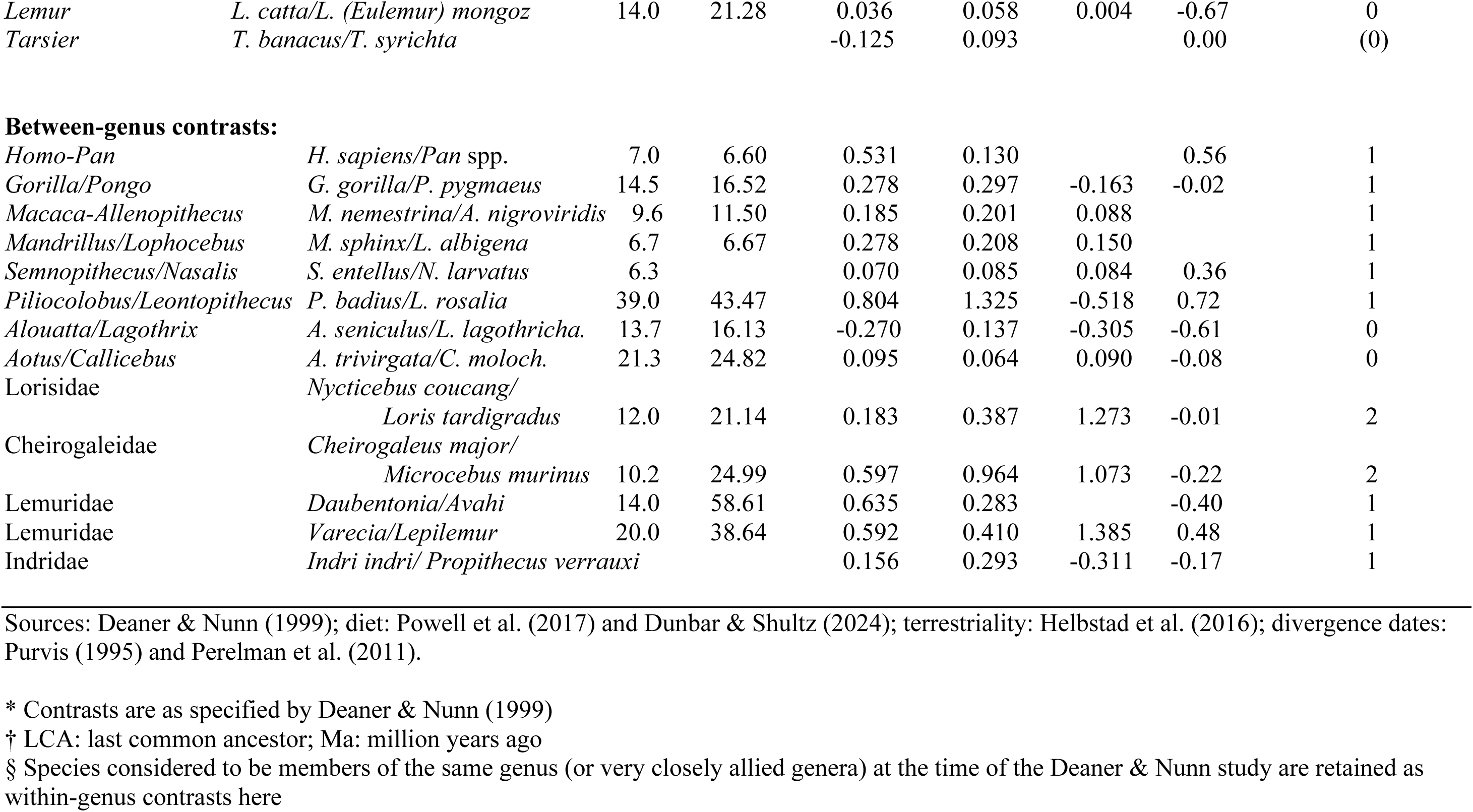
Contrasts in brain and body mass for the sample tip taxa.

**Table S2.** Contrasts for hominins.

| Species | LCA to *Pan*<br>(Ma)* | Contrasts in |  |
| --- | --- | --- | --- |
|  |  | brain mass† | body mass |
| *Australopithecus aethiopicus* | 4.3 | 0.0416 | 0.0110 |
| *Australopithecus afarensis* | 3.2 | 0.0586 | 0.0440 |
| *Australopithecus africanus* | 3.3 | 0.0869 | 0.0440 |
| *Paranthropus boisei* | 4.8 | 0.1170 | 0.0490 |
| *Paranthropus robustus* | 5.0 | 0.1539 | 0.0490 |
| *Homo ergaster* | 5.0 | 0.3180 | 0.2427 |
| *Homo erectus* | 5.4 | 0.4059 | 0.2933 |
| *Homo heidelbergensis* | 6.4 | 0.5102 | 0.3457 |
| *Homo neanderthalensis* | 6.5 | 0.5601 | 0.3528 |
| *H. sapiens* (fossil) | 6.8 | 0.5770 | 0.3316 |

Sources: dates and brain mass: De Miguel & Henneberg (2001); body mass: Will et al. (2017);
*Pan* brain mass from Cofran (2018); *Panm* body mass from Pusey et al. (2005)
\* Million years ago
† ECVs, corrected following Aiello & Dunbar (1993)

